# Processing variables of direct-write, near-field electrospinning impact size and morphology of gelatin fibers

**DOI:** 10.1101/2020.09.17.301804

**Authors:** Zachary G. Davis, Aasim F. Hussain, Matthew B. Fisher

## Abstract

Several biofabrication methods are being investigated to produce scaffolds that can replicate the structure of the extracellular matrix. Direct-write, near-field electrospinning of polymer solutions and melts is one such method which combines fine fiber formation with computer-guided control. Research with such systems has focused primarily on synthetic polymers. To better understand the behavior of biopolymers used for direct-writing, this project investigated changes in fiber morphology, size, and variability caused by varying gelatin and acetic acid concentration, as well as process parameters such as needle gauge and height, stage speed, and interfiber spacing. Increasing gelatin concentration at a constant acetic acid concentration improved fiber morphology from large, planar structures to small, linear fibers with a median of 2.3 μm. Further varying the acetic acid concentration at a constant gelatin concentration did not alter fiber morphology and diameter throughout the range tested. Varying needle gauge and height further improved the median fiber diameter to below 2 μm and variability of the first and third quartiles to within +/-1 μm of the median for the optimal solution combination of gelatin and acetic acid concentrations. Additional adjustment of stage speed did not impact the fiber morphology or diameter. Repeatable interfiber spacings down to 250 μm were shown to be capable with the system. In summary, this study illustrates the optimization of processing parameters for direct-writing of gelatin to produce fibers on the scale of collagen fibers. This system is thus capable of replicating the fibrous structure of musculoskeletal tissues with biologically relevant materials which will provide a durable platform for the analysis of single cell-fiber interactions to help better understand the impact scaffold materials and dimensions have on cell behavior.

## Introduction

The extracellular matrix (ECM) of a tissue, such as fibrous collagen, provides structure as well as biochemical and mechanical cues which can control cellular behavior, such as proliferation and differentiation. Even after tissue decellularization, the ECM retains these functions (1). Not surprisingly then, the field of tissue engineering has implemented a variety of approaches in an attempt to mimic the native ECM of tissues (1–3). Two methods being applied for musculoskeletal fibrous tissue are 3D printing and electrospinning (4–7) due to their ability to fabricate fibrous structures.

There are a variety of different forms of 3D printing, including fused deposition modeling and bioprinting (7–9), which use single fibers to create complex structures. Each method uses software and multi-axis spatial control to deposit individual fibers. The fibers from these systems typically have a minimum range of 100μm-400μm in diameter, which is on the scale of a collagen fascicle (8,10,11). However, some systems, like the freeform reversible embedding of suspended hydrogels, have achieved fiber resolutions down to 20 μm (12,13). Though 3D printing has the potential to replicate the macroscale structure of musculoskeletal fibrous tissues, it currently lacks the ability to achieve the size of smaller fibers and fibrils (14).

Electrospinning uses an electric potential to draw out a stream of fibers from a polymer solution creating a sheet of fibers with diameters that range from nanometers to microns, which approaches the size range of fibrils and fibers (4,10,15). Orientation can be manipulated by using a rotating mandrel or other collection techniques to preferentially align fiber ensembles (16,17). One limitation of electrospinning is that the lack of control over individual fiber placement prevents the creation of complex structures. Some control of individual fibers has been achieved using a spinneret based tunable engineered parameter system combined with a rotating mandrel (18,19). Alternatively, direct-write, near-field electrospinning (DWNFE) combines the software and multi-axis control of 3D printing with the electrohydrodynamic fiber formation of near-field electrospinning to create complex 3D scaffolds (20–22). This approach combines the advantages of the two previous methods allowing for the fabrication of a structure with fibers similar in size to the fibers of collagen (2-20μm) while also meeting the macroscopic requirements needed to replicate musculoskeletal fibrous tissue (20–26). The DWNFE system has even been shown to create 3D stacked fibrous scaffolds with 90-100nm fiber diameters from polyethylene oxide (27).

To date, DWNFE systems have largely been implemented using synthetic polymers, natural biomaterials, or combinations of both (26–32). Synthetic polymers provide mechanical properties similar to musculoskeletal fibrous tissue and are often biodegradable; however, they have limited bioactivity (30,32). Bioactivity within a precisely controllable system is advantageous since the combination of these functions allows for the creation of scaffolds that more actually capture native matrix structure and biomolecular composition to influence cellular behavior (30,31). Two examples of direct-writing with biologics include collagen in an acetic acid solution (32), as well as a gelatin-norbornene (gelNOR) polyethylene oxide (PEO) composite bioinks (31). From this prior work using collagen-based material, it is clear that the acetic acid concentration along with voltage and humidity can alter collagen fiber formation (32). However, other processing parameters, such as needle gauge, needle height, stage speed, and biopolymer concentration, also play important roles in fiber formation in direct-writing approaches (4,22,33,34).

Thus, the objective of this project was to determine the impact of the gelatin and acetic acid concentrations, as well as needle gauge, needle height, and stage speed, on the formation of fibers fabricated through DWNFE. Gelatin is a useful alternative to collagen as it maintains much of the enzymatically active sites, while being more readily available (33,35–37). Gelatin can also be augmented to improve mechanical properties and alter cell function through the addition of various other bioactive molecules (35,38,39). In this work, we show that gelatin and acetic acid concentration have an impact on the viscosity of the solution which in turn impacts the quality and morphology of the gelatin fibers fabricated. By further controlling needle gauge and height, we consistently produced gelatin fibers capable of replicating the collagen fiber structure of musculoskeletal fibrous tissues.

## Materials and Methods

### Formation of Gelatin Solutions

Type A, 300 bloom, porcine gelatin (Sigma-Aldrich Saint Louis, MO, USA) was added to the glacial acetic acid (Fisher Scientific, Fair Lawn, NJ, USA)-diH_2_O. Solutions were mixed at an internal solution temperature of 45°C for 3 days. Initially, the gelatin concentration was varied (450, 500, 525, 550, 600, 625, 650 mg/mL) at a constant 70% acetic acid, then the acetic acid concentration varied (60%, 70%, 80%, or 90% mL/mL) at a constant 625 mg/mL gelatin concentration. Three solutions were made per group. All solutions were stored at 4°C (Fig. S1).

#### Viscosity assessment

The viscosity of solutions was measured on a cup and bob rheometer (MCR 302, Anton Paar, Austria) with a shear rate sweep (0.1-1000 sec^-1^) at 25°C. Briefly, 11 mL of solution was poured into the cup and the bob was lowered so a thin layer of solution covered the top of the bob. A 4-step sequence was used to analyze the rheological response of the solution. To start, a pre-sweep at a shear rate of 1 sec^-1^ was used to remove bubbles and allow for temperature equilibration. Then, the shear rate was ramped up from 0.1 to 1000 sec^-1^. This was followed by a ramp down from 1000 to 0.1 sec^-1^ and finally a second ramp up from 0.1-1000 sec^-1^. The viscosity and shear stress were collected at 40 points throughout each of the shear rate sweep steps. To determine the singular viscosity value for each solution, an average of the viscosity over the linear region for each solution (0.1-100 sec^-1^) was taken.

#### Scaffold fabrication

A custom-built DWNFE system was employed, similar to as previously described (22). Briefly, a desktop, fused deposition 3D printer (Lulzbot Mini, Aleph Objects, USA) was augmented by replacing the tool-head with a custom designed adaptor to hold a 5mL Luer-Loc plastic syringe (Jenson Global, USA) fitted with a blunt metal needle (Jenson Global, USA) (Fig. 1). Pneumatic pressure provided by a Welch pump (Gardner Denver, USA) was provided through the Jenson Global syringe adaptor and regulated for low-pressure control (Dwyer Instruments, USA). Positive voltage (Gamma High Voltage Research, USA) was supplied to the base of the needle to generate the electric potential between the needle and the grounded base underneath the collecting surface of aluminum foil. Prior to printing, the height was tested by lowering the stage until the needle touched the foil. This point was designated as the origin of the vertical axis. MATLAB was used to generate a G-code grid patterned tool path for the 3D printer. The solution was poured into the 5mL syringe and mounted on the location of the tool head. A voltage ranging from 1 kV to 6 kV was applied as was a pressure between 0 and 0.4 psi (Table S1). Scaffold dimensions were 20 mm by 20 mm and 2 layers tall with an initial fiber spacing of 1mm. Printing occurred at an ambient humidity of 48-54%. Three scaffolds were made from each of the 3 solutions per group with each solution being used on a different day. In total, 9 scaffolds were fabricated for each group.

**Figure 1.**
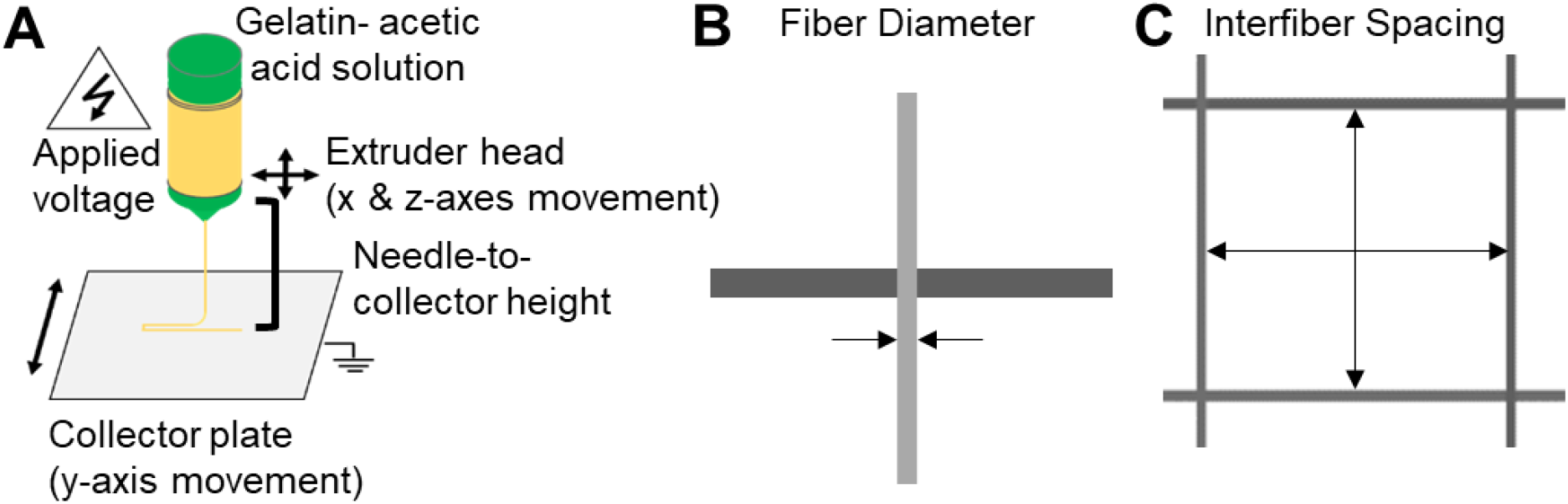
A) Schematic of the DWNFE process. B) Top-down view of fibers with arrows indicating fiber diameter. C) Top-down view of fibers with arrows indicating interfiber spacing.

The process parameters of stage speed (15, 30, 45, 60, 75, or 90 mm/s), needle height (1, 2, or 4 mm), and needle gauge (20 G (ID=0.610 mm), 22 G (ID=0.406 mm), and 25 G (ID=0.254 mm)) were also investigated for the 70% acetic acid, 625 mg/mL gelatin solution. Needle height was varied for each needle gauge at a stage speed of 45 mm/s. For stage speed, a 22 G needle and 1mm height was used. Finally, fiber spacing (1000, 500, 250, 100, and 50 μm) was varied with constant stage speed of 45 mm/s, 22 G needle, and 1 mm height. The voltage and pressure were adjusted for optimal fiber formation for each group. Square samples (1 cm × 1 cm) were cut from 3 scaffolds per group per day on 3 different days for further analysis.

#### Fiber and spacing analysis

Scaffold samples were sputter coated for ∼1 minute to create a conductive surface for imaging via scanning electron microscopy (SEM). Two different SEM devices were used due to machine availability. A S3200N SEM (Hitachi, Japan) was used at X40, X70, and X1000 with an accelerating voltage of 5kV for studies of concentration and stage speed. A JCM 7000 SEM (JEOL, USA) was used at X40, X70, and X1000 with a 5kV accelerating voltage was used for studies of spacing, needle gauge, and needle height. SEM images were analyzed via ImageJ (NIH). Fiber diameter and interfiber spacing were determined for each group by measuring 6 fiber diameters and 6 interfiber spacings for 9 specimens per group (Fig. 1). This gave 54 fibers and 54 interfiber spacings measurements per group.

### Statistical Analysis

Normality was determined via the D’Agostino-Pearson test. Outliers were then determined by the ROUT test with Q= 0.1%. Outliers were then removed from the data groups before running the nonparametric Mann-Whitney t-tests (α=0.05) to determine statistically significant differences between groups for the fiber study comparisons. Desired thresholds for fiber diameter were set to <2 μm fiber diameter and quartiles within +/-1μm delta from the median for each group. For interfiber spacing, a one sample Wilcoxon test was used to compare the percent error to the ideal error value (zero) for each spacing (α=0.05). A threshold for the quartiles of +/-20% error relative to the intended value was set for the spacing.

## Results

### Gelatin concentration

The viscosity of the gelatin solution increased with increasing gelatin concentration (Fig. 2B, Table S2). Values were steady from 450 to 500 mg/mL with median viscosities of 1.3 and 1.0 Pa*s respectively. This was followed by an increase in median viscosity at 525 mg/mL (2.1 Pa*s) and 550 mg/mL (2.5 Pa*s) before a 3-fold jump for the 600 and 625 mg/mL groups (6.6 and 6.4 Pa*s, respectively). Finally, for the 650 mg/mL group, the viscosity increased again to 8.0 Pa*s but had greater variability than seen in the other groups.

**Figure 2.**
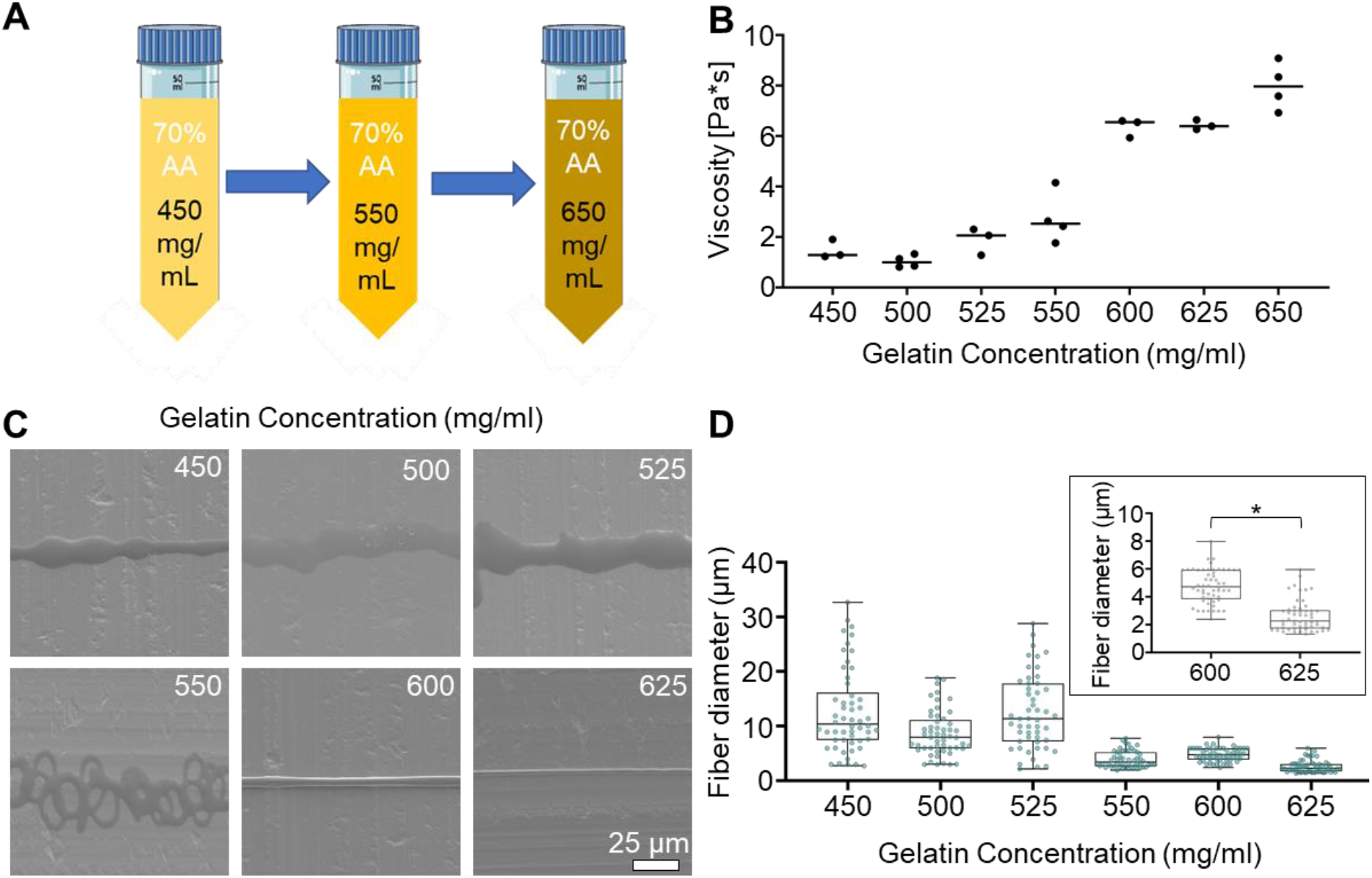
Varying gelatin concentration altered the morphology of the produced fibers. A) Schematic of range of gelatin concentration tested at a constant 70% acetic acid concentration. B) Viscosity of gelatin solutions increased with increasing gelatin concentration (n=3-4 per concentration; median represented with a bar). C) Fiber morphology improves to form continuous, linear fibers above 600 mg/mL gelatin concentrations (scale bar = 25 μm). Note: 650mg/mL was not direct-written due to the variability in the solutions. D) Fiber diameter decreases with increasing gelatin concentration. Statistics were not performed for the lower gelatin concentrations due to poor fiber morphology. The insert is zoomed in on the 600 and 625 mg/mL from panel D (data are presented as median and IQR for n=51-54 per concentration; * indicates a statistically significant difference between the groups).

Fiber morphology and diameter were dependent on gelatin concentration (Fig. 2C-D, S2, Table S3). The lower concentration groups, 450-525 mg/mL, had an inconsistent, broad, and planar morphology. There was little change in the morphology between these lower concentration groups. The fiber diameter for these groups was large with a high variability; for example, the 450 mg/mL group had a median value of 10.4 μm and IQR 7.4-16.15 μm. For the 550 mg/mL group, thinner and more consistent, though curled, lines were produced, with a median fiber diameter of 3.4 μm and IQR 2.6-5.2 μm, though these fibers still had a broad, flat morphology. At 600mg/mL, a thin, continuous fiber morphology with a median fiber diameter of 4.7 μm and an IQR of 3.8-5.9 μm was observed with a 3D cylindrical shape instead of the flat lines seen at lower concentrations. At 625 mg/mL, the same thin continuous fiber morphology was seen but with a smaller median fiber diameter of 2.3 μm and IQR of 1.7-3.0 μm. The cylindrical fiber structure was verified through cross-sectional SEM (Fig. S3). Between the two groups that provide an appropriate fiber morphology (600 and 625 mg/mL), there was a statistically significant difference in fiber diameter (p<0.0001). Overall, fiber diameter decreased with increasing gelatin concentration, while the fiber morphology changed from inconsistent flat structures to well-formed cylindrical fibers.

### Acetic acid concentration

Using a constant gelatin concentration of 625 mg/mL, the viscosity of the gelatin solutions increased with increasing acetic acid concentration (Fig. 3B, Table S2). Similar values were obtained for the 60% and 70% acetic acid groups (5.9 Pa*s and 6.4 Pa*s, respectively). However, when the concentration was further increased, the mean viscosity approximately doubled to 10.1 Pa*s at 80% acetic acid and doubled again to 22.9 Pa*s at 90% acetic acid. The viscosity of the 90% acetic acid group was highly variable and not used in subsequent analyses.

**Figure 3.**
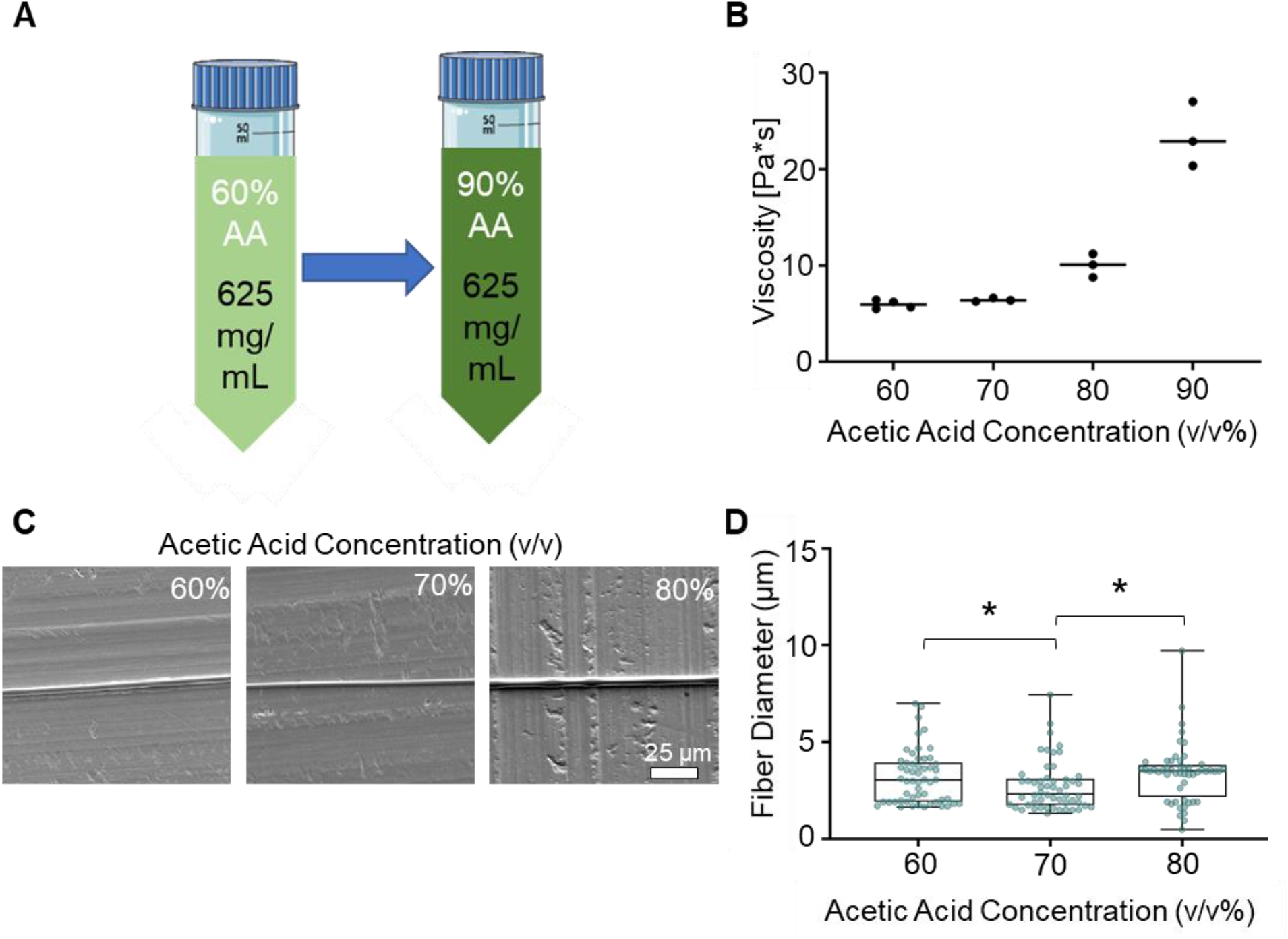
Varying the acetic acid (AA) concentration altered the morphology of the produced fibers. A) Schematic of range of acetic acid concentration with a constant 625mg/ml gelatin concentration. B) Viscosity of gelatin solutions increased with increasing AA concentration (n=3-4 per concentration; median represented with a bar). C) No change in morphology was seen when the acetic acid concentration was changed (scale bar = 25 μm). Note: 90% AA was not direct-written due to variability in the viscosity. D) The smallest and most consistent fibers were formed at 70% AA (data are presented as median and IQR for n=52-54 per concentration * indicates a statistically significant different between the groups).

Fiber morphology did not change with varied acetic acid concentration (Fig. 3C-D, Table S3). The fibers formed at 60, 70 and 80% acetic acid were all linear, continuous fibers with median diameters of 3.0, 2.3, and 3.5 μm and IQRs of 1.9-3.9, 1.7-3.0, and 2.1-3.8 μm respectively (Fig. 3D). A statistically significant difference in fiber diameter was found between the 70% acetic acid group and the 60 and 80% acetic acid groups (p=0.02 for 60 vs. 70%; p=0.001 for 70 vs. 80%); however, the magnitudes of these median differences (0.7 μm and 1.2 μm for 60 vs. 70% and 80 vs. 70%, respectively) were small.

### Needle gauge and height

Using a solution of 625 mg/mL gelatin and 70% acetic acid, the impact of needle gauge and height was assessed, and both variables affected fiber diameter (Fig. 4A, Table S4). Thresholds of successful fiber production were set at a maximum of 2 μm for fiber diameter and +/-1 μm from the median for variability. Initially, the 20 G needle size was tested at 1-, 2-, and 4-mm heights. A 23% increase in median diameter was seen between 1 mm and 2 mm, while between 2 mm and 4 mm a 64% increase was seen. The median fiber diameters at 1 and 2 mm were below the threshold; however, the fiber diameter was greater than 2 μm at 4 mm. The variability as measured by the delta from the median had first and third quartiles within less than the +/- 1 μm of the median at all heights (Fig. 4C, Table S5).

**Figure 4.**
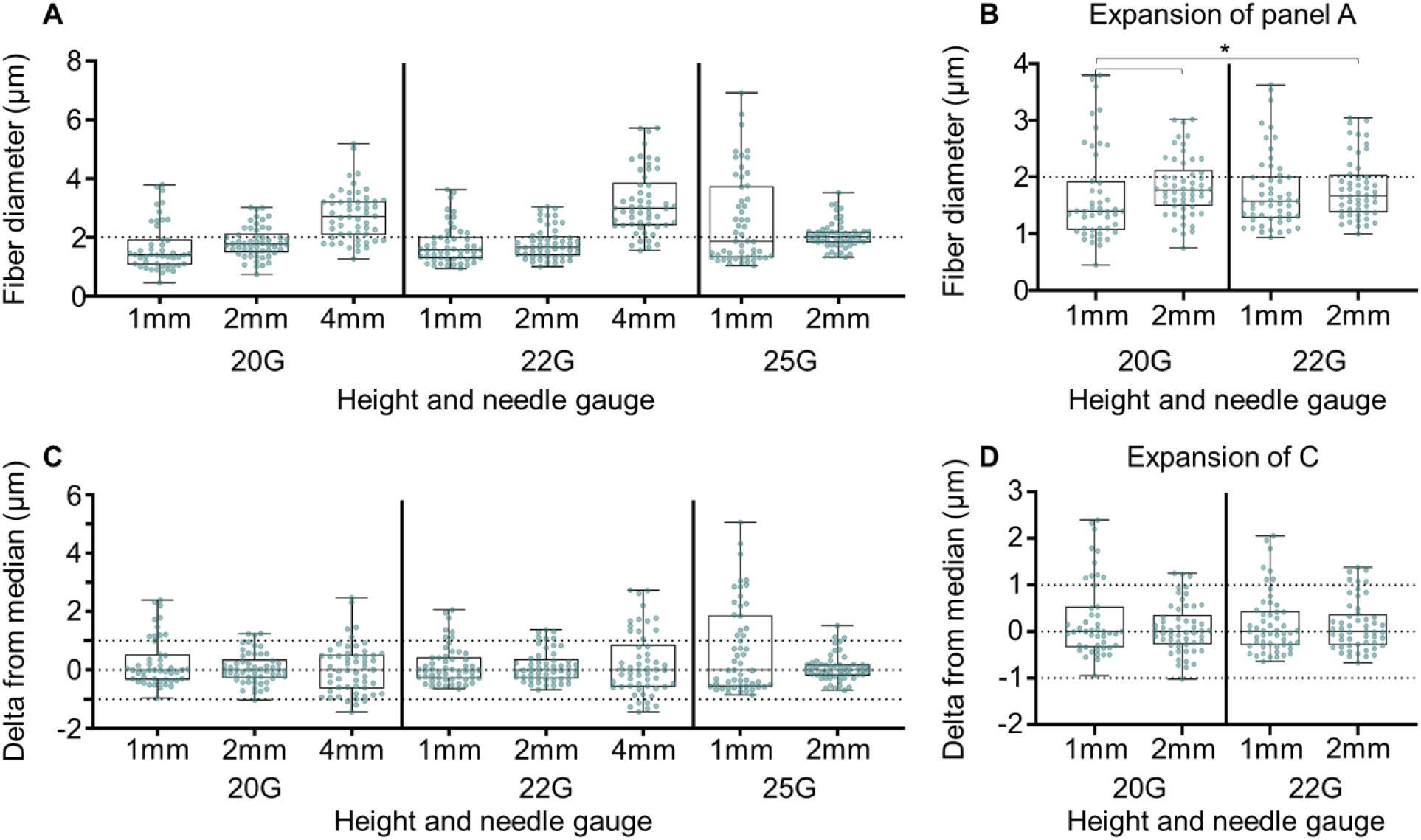
A) Varying the needle gauge and height further impacted the fiber diameter of the 625 mg/mL gelatin and 70% acetic acid solution. (dashed line indicates 2 μm threshold). Note: 4 mm 25 G was not able to be direct-written. B) Expansion of the 1 mm and 2 mm 20 G, and 1 mm and 2 mm 22 G groups from panel A. C) Delta of the measured fiber diameters from the median for each group (dashed line indicates +/-1 μm threshold). D) Expansion of the 1 mm and 2 mm heights with the 20 G and 22 G needles groups from panel C (data are presented as median and IQR for n=47-54 per group; * indicates a statistically significant different between the groups).

Based on these results, we decided to implement a smaller needle size (22 G, Fig. 4A, Table S4). The same increase in median fiber diameter with height was seen within the 22 G groups with a 6% difference between the 1 mm and 2 mm groups and a 56% difference between the 2 mm and 4 mm groups. The median fiber diameters at 1 and 2 mm were below the 2 μm threshold. The variability of the first and third quartiles from the median for all heights were within the +/-1 μm threshold (Fig. 4C, Table S5). Relative to the 20 G groups, fiber sizes were similar with median fiber sizes using a 22 G needle within 10-18% controlling for needle height. A small decrease in median diameter from 20 G to 22 G (18% difference) was noted at 2 mm.

To test this further, 25 G needles were used. The 25 G needles produced large fibers at 1 mm compared to the 20 G (29% difference) and 22 G (17% difference) groups with a median diameter of 1.87 μm which was still within the 2 μm threshold (Fig. 4A, Table S4). However, the variability was also beyond the +/-1 μm threshold (Fig. 4C, Table S5). At 2 mm needle height, the median fiber diameter increased to 2.01 μm, but the variability decreased to within the threshold showing more consistent fiber deposition. When the height was increased to 4 mm, the 25 G needle was not able to form fibers. Across all tests, the groups with median fiber diameters below the median threshold of 2 μm and within the +/- 1 μm threshold were compared statistically (Fig. 4B&D). Significant differences were noted between 1 and 2 mm 20 G (p=0.01) and 1 mm 20 G and 2 mm 22 G (p=0.02).

### Stage speed

Using a solution of 625 mg/mL gelatin and 70% acetic acid concentration with the 22 G needle at 1 mm height, stage speed had no additional effect on the morphology of gelatin fibers (Fig. 5A, Table S6). Continuous fibers were produced at each speed. Fiber diameter was not affected by stage speed with statistically significant differences seen between the 15 mm/s group and the 60, 75 and 90 mm/s groups (p=0.001 for 15 mm/s vs. 60 mm/s; p=0.003 for 15 mm/s vs. 75 mm/s; p<0.001 for 15 mm/s vs. 90 mm/s) as well as between 30 mm/s and 90 mm/s (p=0.046)(Fig. 5B).

**Figure 5.**
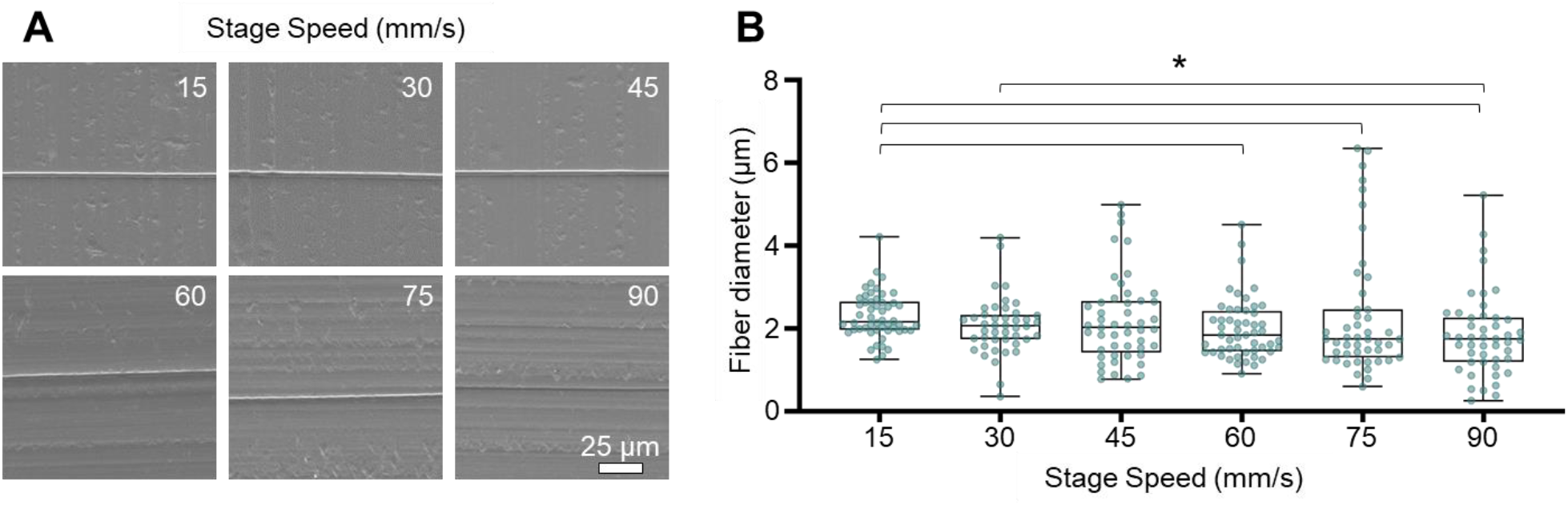
A) Varying stage speed did not alter fiber morphology for the 625mg/mL gelatin 70% acetic acid solution (scale bar = 25 μm). B) Varying stage speed did not alter fiber diameter, but a slight decrease in diameter as the speed is increased is visible (data are presented as median and IQR for n=44-52 per group; * indicates a statistically significant different between the groups).

### Spacing

Using optimized parameters, the accuracy of our system to achieve a theoretical spacing as a percent error was analyzed (Fig. 6, S4, Table S7). High variability was seen when attempting to create a scaffold with 50 μm spacing with a median percent error of 70.3% and IQR of 41.6 to 92.2% (Fig. 6B). The median percent error between the measured interfiber spacings for the 250, 500, and 1000 μm and the theoretical spacing were not statistically significantly different from 0 (250 μm p=0.13; 500 μm p=0.27, and 1000 μm p=0.48) (Fig. 6C). The 100 μm group, however, was statistically different with median percent errors of 5.2% IQR of -5.7 to 22.6%. The medians and IQRs for all groups aside from the 50 and 100 μm groups were within the 20% error threshold.

**Figure 6.**
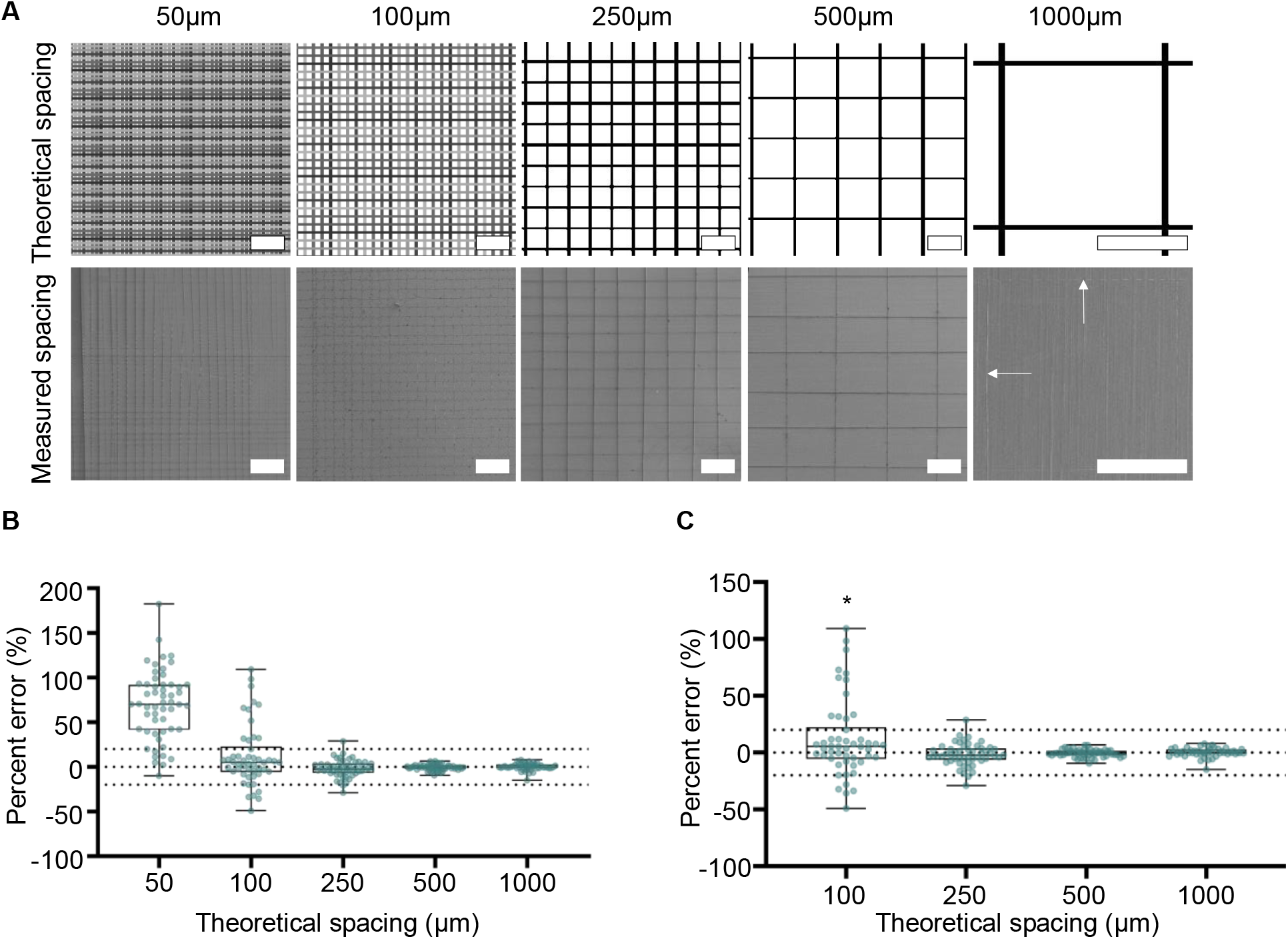
Interfiber spacing of 625mg/mL gelatin 70% acetic acid solution at a constant 45mm/s stage speed. A) The MATLAB generated 2-layer grid patterns for each spacing and representative SEM images at X40 for 50-500 μm and X70 for 1000 μm (scale bar = 500 μm). Arrows indicate fibers in the 1000 μm spacing image. B) The measured spacings for 1000 μm, 500 μm, and 250 μm were not different from the theoretical spacing. C) Is an expansion of B without the 50 μm group (data are presented as median and IQR for n=48-54 per spacing; dashed lines indicate the +/- 20% error threshold; * indicates a statistically significant different from 0).

## Discussion

In this study, we have shown that gelatin and acetic concentration, as well as needle gauge and height, impact gelatin fiber formation during direct-write, near-field electrospinning. Once these parameters were optimized, additional variation of stage speed did not have a major impact. Our results show the ability to produce scaffolds with consistent fiber morphology and controllable diameters less than 2 μm (+/-1 μm) along with spacings from 250 to 1000 μm with less than 20% error.

Gelatin concentration affected the viscosity since increasing the concentration increased the viscosity, as also shown by Erencia et al (33). This viscosity change led to both fiber diameter and fiber morphology changes with lower concentrations generating planar structures and higher concentrations, distinctive linear fibers. This is corroborated by Castilho et al who also showed an inverse correlation between concentration and fiber diameter (31). The results show that a viscosity of 6-7 Pa*s produces linear fibers at 50% humidity. This humidity was higher than was shown to produce consistent fibers by Alexander et al. (32) for collagen, but this difference may be due to altering the gelatin concentration as well as differences in gelatin versus collagen solutions. The optimal fibers were determined to be the ones produced at 625 mg/mL gelatin with a median value of 2.3 μm and IQR of 1.7-3.0 μm. This median is similar to the fiber diameters (∼2 μm) and structures seen by collagen and gelNOR fibers in prior work as well as matching the extreme low end of typical PCL MEW fibers (2 μm) (31,32,40). They, however, do not reach the 92 nm fiber sizes see with prior work using polyethylene oxide solution (27).

Acetic acid was seen to have a similar effect on the viscosity of gelatin-acetic acid solutions with increasing concentration increasing the viscosity. This corroborates the findings of prior work showing that increasing acetic acid concentrations from 25 to 75 (v/v)% increased the viscosity of the solution (33). Groups with 60, 70, and 80% acetic acid were able to produce linear and consistent fibers, and even though a statistical difference was found between the groups, it is not clear what a 1 μm median fiber difference means in terms of utility in tissue engineering applications. From this, we can conclude that each of these concentrations is viable for use, but we applied the 70% acetic acid for future studies.

Varying printing parameters, such as the needle gauge, needle-to-collector height, and stage speed affects both the fiber diameter and variability of produced fibers (4,20,22). Brown et al. have previously shown that decreasing needle size decreases fiber diameter; while Warren et al. previously showed changing the needle height can affect fiber deposition for fabricating PCL melt written scaffolds (17,19). Fuh et al. previously tested the impact of needle-to-collector height for chitosan-poly(ethylene oxide) hybrid solution and showed a decrease in fiber diameter with increasing distance (34). The effect of needle gauge on solution direct-writing of biopolymers was not known as only 2 gauges (22 and 27 G) have been used within different previous studies (23,26,28). To test the impact of different needle gauges, we tested 20 G (ID=0.610mm), 22 G (ID=0.406 mm), and 25 G (ID=0.254 mm). To test the impact of needle height, each needle gauges was sampled at three different heights (1 mm, 2 mm, and 4 mm). For both 20 G and 22 G needles, an increase in fiber diameter was seen with increasing height, while the variability was not significantly affected. For both gauges, the 4 mm height produced the largest fibers with medians above the 2 μm threshold, with the greatest variability. When the 25 G needle was used, the needle was more likely to clog than the previous needles which may explain the variability seen for the 1 mm height. Additionally, fibers were not able to form at 4 mm because of the electric potential discharge that occurred. This was due to an inability to maintain a consistent amount of material at the tip without the gelatin drying which caused a spark whenever the flow of solution was disrupted. From these results, we determined that the 20 G or 22 G at 1 mm or 2 mm height was optimal.

Stage speed has previously been shown to be a factor of interest for MEW of PCL by significantly decreasing fiber diameter from 80 μm to 25 μm by increasing stage speed from 1-5 mm/s (22). However, in this study, no significant change in fiber morphology or diameter was detected. A slight, statistically significant decrease was observed between the 15 mm/s and 75 and 90 mm/s groups in the current work, but the decrease is unlikely to be experimentally significant. Speed may impact melt writing but not solution writing because of viscosity differences, as the molten polymer is significantly more viscous. This may cause mechanical drawing of the fiber which would decrease the diameter as the speed was increased. Another possible cause for the different impact seen is that the conductivity of the polymer may affect the speed of the polymer jet (24). Other parameters such as voltage and pressure have been shown to also impact fiber formation, but in this study, we assumed they were dependent upon the parameters tested here (20,28,32).

Finally, we needed to test the limits of the interfiber spacing capability of the system. Interfiber spacing plays a key role in regulating cell function (28). We tested a range of spacings from 50-1000 μm and showed 250-1000 μm spacings are capable with less than a 20% error. This does not reach the 10-20 μm spacings of Fuh et al. because of the stepper motor size limitations of our system (28). However, these spacings will be valuable for analyzing single cell-fiber interactions.

This study has shown the application and optimization of gelatin-acetic acid solutions for DWNFE; however, some limitations exist. For one, not all possible combinations were tested. This means there may be another combination of acetic acid and gelatin concentration, and process parameters that also produces functional fibers. The conclusions on the optimal groups from this study are valuable for our future experiments and have been shown to be reproducible through fabrication with different solutions and on different days. Another limiting factor was humidity which has been shown to have an impact on printing and was not fully controlled in these experiments (32). Thus, further experimentation into the impact of humidity and optimizing concentrations and parameters at different levels of humidity may be helpful.

These results add to the growing body of literature using electrowriting to engineer tissues and study cell-biomaterial interactions (9,18,30–32,40–42). Importantly, these gelatin fibers are on the diameter range of collagen fibers which is the domain where the fibroblast would reside, while also providing a more natural biomaterial than synthetic polymers (10). The control over fiber diameter and location of natural polymers will allow exploration of cell-ECM interactions critical to the function of many tissues, including fibrous musculoskeletal soft tissues.

## Conclusions

In conclusion, increasing gelatin and acetic acid concentrations increases viscosity which, to a point, improves fiber morphology and decreases diameter to around a 2 μm threshold. By altering the process parameters, the diameter was reduced to below the 2 μm threshold and the variability was able to be decreased to within the +/-1 μm threshold. Along with showing the system’s ability to produce consistent fibers this study also found the interfiber spacing limit of the system to be 250 μm. The control over fiber morphology and diameter we have demonstrated will allow us to create unique natural scaffolds for tissue engineering.

## Acknowledgements

This work was supported in part by the North Carolina State University Game-Changing Research Incentive Program (GRIP) and was performed in part at the Analytical Instrumentation Facility (AIF) at North Carolina State University, which is supported by the State of North Carolina and the National Science Foundation (award number ECCS-1542015). The AIF is a member of the North Carolina Research Triangle Nanotechnology Network (RTNN), a site in the National Nanotechnology Coordinated Infrastructure (NNCI). This material is based upon work supported by the National Science Foundation Graduate Research Fellowship Program under Grant No. DGE-1746939. Any opinions, findings, and conclusions or recommendations expressed in this material are those of the author(s) and do not necessarily reflect the views of the National Science Foundation. Work was also performed at the North Carolina State University Food Rheology Laboratory with support from Chris Pernell. Special thanks to Carina Iboaya for her work on sample production.

