## Supplemental materials for "Processing variables of direct-write, near-field electrospinning impact size and morphology of gelatin fibers"

Table S1: Gelatin and acetic acid concentrations and processing parameters used during the experiments.

| Key variable | Group concentrations | | Process Parameters | | | | |
|---|---|---|---|---|---|---|---|
| | Acetic acid (v/v%) | Gelatin (mg/mL) | Pressure (psi) | Stage speed (mm/s) | Height (mm) | Needle gauge (G) | Voltage (kV) |
| **Gelatin concentration (Fig. 2)** | 70 | 450-650 | 0 - 0.1 | 45 | 2 | 22 | 3-6 |
| **Acetic acid concentration (Fig. 3)** | 60-90 | 625 | 0.1 - 0.4 | 45 | 1 | 22 | 3 |
| **Process parameters (Fig. 4&5)** | 70 | 625 | 0 - 0.1 | 15-90 | 1, 2, 4 | 20, 22, 25 | 3 |

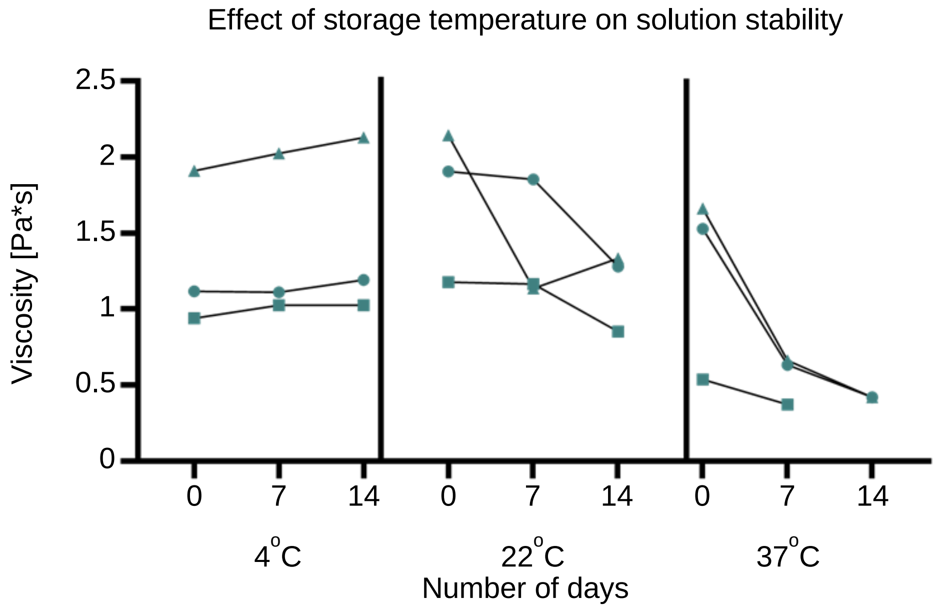


Figure S1: Viscosity of 70% acetic acid, 45mg/mL gelatin solution stored at different temperatures over 2 weeks. At 4°C the solutions maintained the starting viscosity or slightly increased, while at 22°C and 37°C the viscosity decreased with time. The black line indicates the median for n=3 solutions.

Table S2: Summary of solution viscosities.

| | Gelatin concentration (mg/mL) | | | | | | | Acetic acid concentration (v/v%) | | | |
|---|---|---|---|---|---|---|---|---|---|---|---|
| | 450 | 500 | 525 | 550 | 600 | 625 | 650 | 60 | 70 | 80 | 90 |
| **Median Viscosity (Pa*s)** | 1.3 | 1.0 | 2.1 | 2.5 | 6.6 | 6.4 | 8.0 | 5.9 | 6.4 | 10.1 | 22.9 |

Table S3: Summary statistics for fiber diameters from altering gelatin and acetic acid concentration. *Statistically significant from 625mg/mL for gelatin concentration and 70%AA for acetic acid concentration.

| | Gelatin concentration (mg/mL) | | | | | | Acetic acid concentration (v/v%) | | |
|---|---|---|---|---|---|---|---|---|---|
| | **450** | **500** | **525** | **550** | **600*** | **625** | **60*** | **70** | **80*** |
| **Median (μm)** | 10.4 | 8.0 | 11.4 | 3.4 | 4.7 | 2.3 | 3.0 | 2.3 | 3.5 |
| **Q1 (μm)** | 7.4 | 5.9 | 7.2 | 2.6 | 3.8 | 1.7 | 1.9 | 1.7 | 2.1 |
| **Q3 (μm)** | 16.15 | 11.1 | 17.8 | 5.2 | 5.9 | 3.0 | 3.9 | 3.0 | 3.8 |
| **IQR (μm)** | 8.75 | 5.2 | 10.6 | 2.6 | 2.1 | 1.3 | 2.0 | 1.3 | 1.7 |

Table S4: Summary statistics for fiber diameters from altering needle gauge and needle height. *Indicates within the thresholds for median fiber diameter (2um).

| **Needle Gauge** | **20G** | | | **22G** | | | **25G** | |
|---|---|---|---|---|---|---|---|---|
| **Height** | **1mm*** | **2mm*** | **4mm** | **1mm*** | **2mm*** | **4mm** | **1mm*** | **2mm** |
| **Median (μm)** | 1.40 | 1.77 | 2.71 | 1.57 | 1.67 | 2.98 | 1.87 | 2.01 |
| **Q1 (μm)** | 1.07 | 1.49 | 2.08 | 1.28 | 1.38 | 2.41 | 1.31 | 1.82 |
| **Q2 (μm)** | 1.93 | 2.12 | 3.23 | 2.02 | 2.04 | 3.85 | 3.74 | 2.20 |
| **IQR (μm)** | 0.86 | 0.63 | 1.15 | 0.74 | 0.66 | 1.44 | 2.43 | 0.38 |

Table S5: Summary of the delta from median values for the needle gauge and height.* Indicates within the threshold for median variability (within +/-1um).

| **Needle Gauge** | **20G** | | | **22G** | | | **25G** | |
|---|---|---|---|---|---|---|---|---|
| **Height** | **1mm*** | **2mm*** | **4mm*** | **1mm*** | **2mm*** | **4mm*** | **1mm** | **2mm*** |
| **Median (μm)** | 0.00 | 0.00 | 0.00 | 0.00 | 0.00 | 0.00 | 0.00 | 0.00 |
| **Q1 (μm)** | -0.33 | -0.28 | -0.63 | -0.29 | -0.29 | -0.57 | -0.56 | -0.19 |
| **Q2 (μm)** | 0.53 | 0.36 | 0.52 | 0.44 | 0.37 | 0.87 | 1.87 | 0.19 |
| **IQR (μm)** | 0.86 | 0.64 | 1.15 | 0.73 | 0.66 | 1.44 | 2.43 | 0.38 |

Table S6: Summary statistics for fiber diameters from altering stage speed. *Statistically significantly different from 15mm/s.

| | **Stage Speed (mm/s)** | | | | | |
|---|---|---|---|---|---|---|
| | **15** | **30** | **45** | **60*** | **75*** | **90*** |
| **Median (μm)** | 2.2 | 2.1 | 2.0 | 1.8 | 1.7 | 1.8 |
| **Q1 (μm)** | 2.0 | 1.7 | 1.4 | 1.5 | 1.3 | 1.2 |
| **Q3 (μm)** | 2.6 | 2.3 | 2.7 | 2.4 | 2.5 | 2.3 |
| **IQR (μm)** | 0.6 | 0.6 | 1.3 | 0.9 | 1.2 | 1.1 |

Table S7: Summary of interfiber spacing percent errors. *Statistically significantly different from 0.

| | Theoretical interfiber spacing | | | | |
|---|---|---|---|---|---|
| | **1000** | **500** | **250** | **100*** | **50*** |
| **Median (%)** | 0.0 | -0.4 | -2.6 | 5.2 | 70.3 |
| **Q1 (%)** | -1.2 | -2.5 | -6.6 | -5.7 | 41.6 |
| **Q3 (%)** | 2.7 | 1.6 | 3.6 | 22.6 | 92.2 |
| **IQR (%)** | 3.9 | 4.1 | 10.2 | 28.3 | 50.6 |

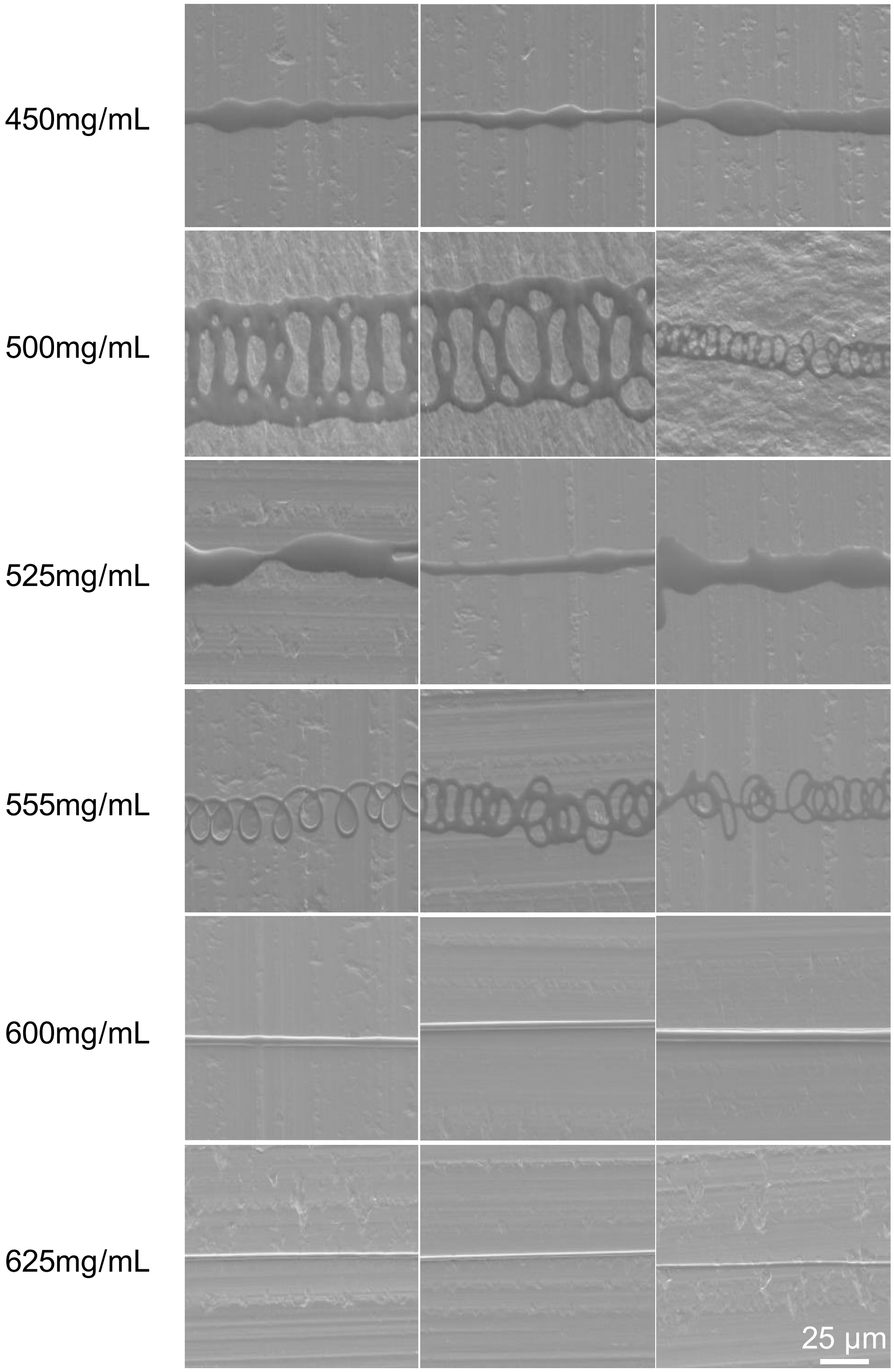


Figure S2: SEM images at X1000 magnification Collection of fibers from different scaffolds produced at different gelatin concentrations (scale bar 25 µm).

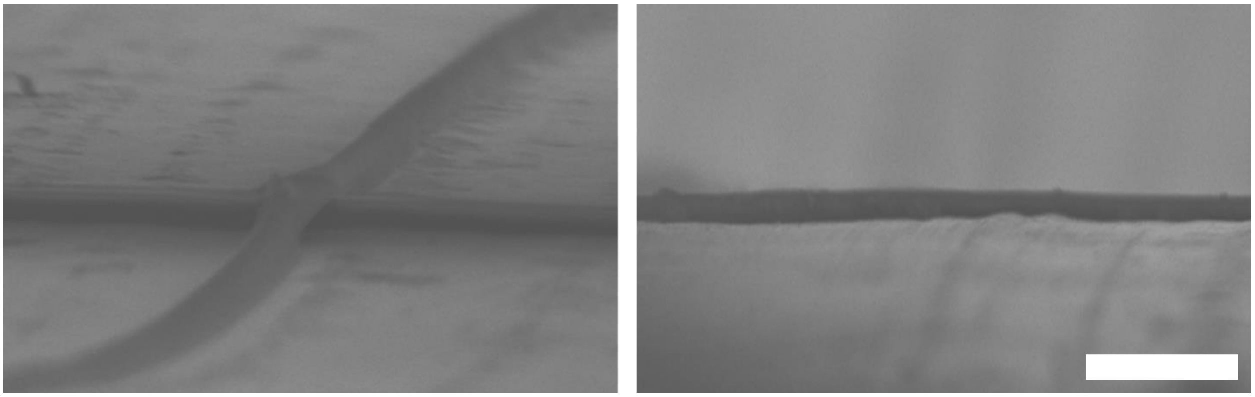

Figure S3: SEM images at X1500 of the sideview of representative scaffolds (scale bar 10 μm).

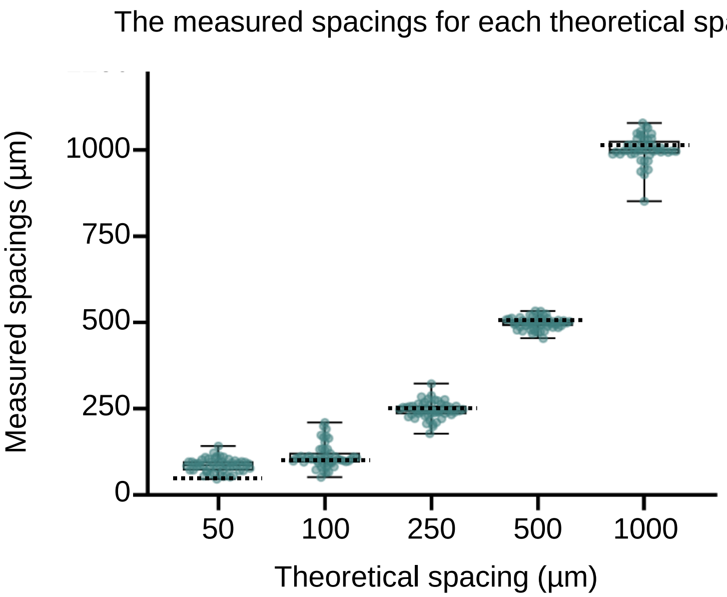


Figure S4: Measured values of the interfiber spacing. The dashed lines represent the theoretical spacing for each group.
